# BiomeHorizon: visualizing microbiome time series data in R

**DOI:** 10.1101/2021.08.29.458140

**Authors:** Isaac Fink, Richard J. Abdill, Ran Blekhman, Laura Grieneisen

## Abstract

**Summary:** A key aspect of microbiome research is analysis of longitudinal dynamics using time series data. A method to visualize both the proportional and absolute change in the abundance of multiple taxa across multiple subjects over time is needed. We developed *BiomeHorizon*, an open-source R package that visualizes longitudinal compositional microbiome data using horizon plots.

**Availability and Implementation:** BiomeHorizon is available at https://github.com/blekhmanlab/biomehorizon/ and released under the MIT license. A guide with step-by-step instructions for using the package is provided at https://blekhmanlab.github.io/biomehorizon/. The guide also provides code to reproduce all plots in this manuscript.

**Supplementary information:** None

## Introduction

Despite playing a key role in the health of their hosts (McKenney *et al*., 2018; Costello *et al*., 2012), host-associated microbial communities demonstrate considerable variation both between hosts and within an individual host over time (Flores *et al*., 2014; Human Microbiome Project Consortium, 2012; Johnson *et al*., 2019). To determine drivers of this temporal variation, and to link such variation to specific host health outcomes, recent work has focused on collecting time series microbiome samples from individual hosts. However, host-associated microbiome data is compositionally complex, with thousands of microbial taxa present at any given time point, and visualizing these longitudinal data is challenging. Traditional methods of longitudinal microbiome visualization use a stream or line graph with a different color for each microbe (Johnson *et al*., 2019; David *et al*., 2014; Turroni *et al*., 2017; Baksi *et al*., 2018). While this is valuable for tracking a single microbe in a single host, it becomes visually difficult to distinguish broader trends among several microbes. This is especially true given that large proportional changes in microbes with low abundances (i.e. mean relative abundance <0.5%) are dwarfed by highly abundant microbes (i.e. mean abundance >25%). Further, line graphs do not facilitate comparing microbial trends at the same time points across different hosts. Among other software tools for longitudinal microbiome analysis, few allow for automated visualization of proportional changes in fluctuations for multiple microbes, or within multiple hosts, over time.

## Materials and methods

Here, we present *BiomeHorizon*, a *ggplot2*-compatible R package that provides a compact way of visualizing the longitudinal dynamics of multiple microbes in parallel (Wickham, 2016). *BiomeHorizon* was developed with two of the most common microbiome study designs in mind: human health experiments, such as dietary or medical intervention studies, where the microbiome is sampled from all subjects at set time points to compare microbiome health outcomes over time; and observational wildlife studies, where samples may be collected at irregular time intervals (e.g., opportunistically when defecation is observed) and/or with large time gaps.

*BiomeHorizon* generates horizon plots, a chart where the X-axis is a time series and the Y-axis starts from the “horizon” (or “origin”; often the median value of a variable across all time points) and creates area charts whose Y-axis distribution represents the distance of the variable from the origin at a given time point (**Fig. 1A**) (Heer *et al*., 2009). Colored bands are used to represent *n*-tiles from the origin, with two different color families representing positive or negative values. The compactness of horizon plots facilitates easy pattern-recognition and comparison between numerous time series, revealing unique insights into longitudinal data. Specifically, they allow visual identification of sustained versus temporary change in microbe(s), which is valuable for modeling stability and disturbance (David et al. 2014). For example, it is easy to detect “comovement” or “periodicity”, while also comparing amplitude. Comovement can be valuable for identifying microbes with related functions (e.g. if they increase at the same proportion over the same timescale), while periodicity might reveal links between environmental or experimental factors and microbial populations. All this functionality is made customizable by *BiomeHorizon*, which provides the flexibility to emphasize specific dimensions of the data and to adapt to unique study designs.

**Figure 1.**
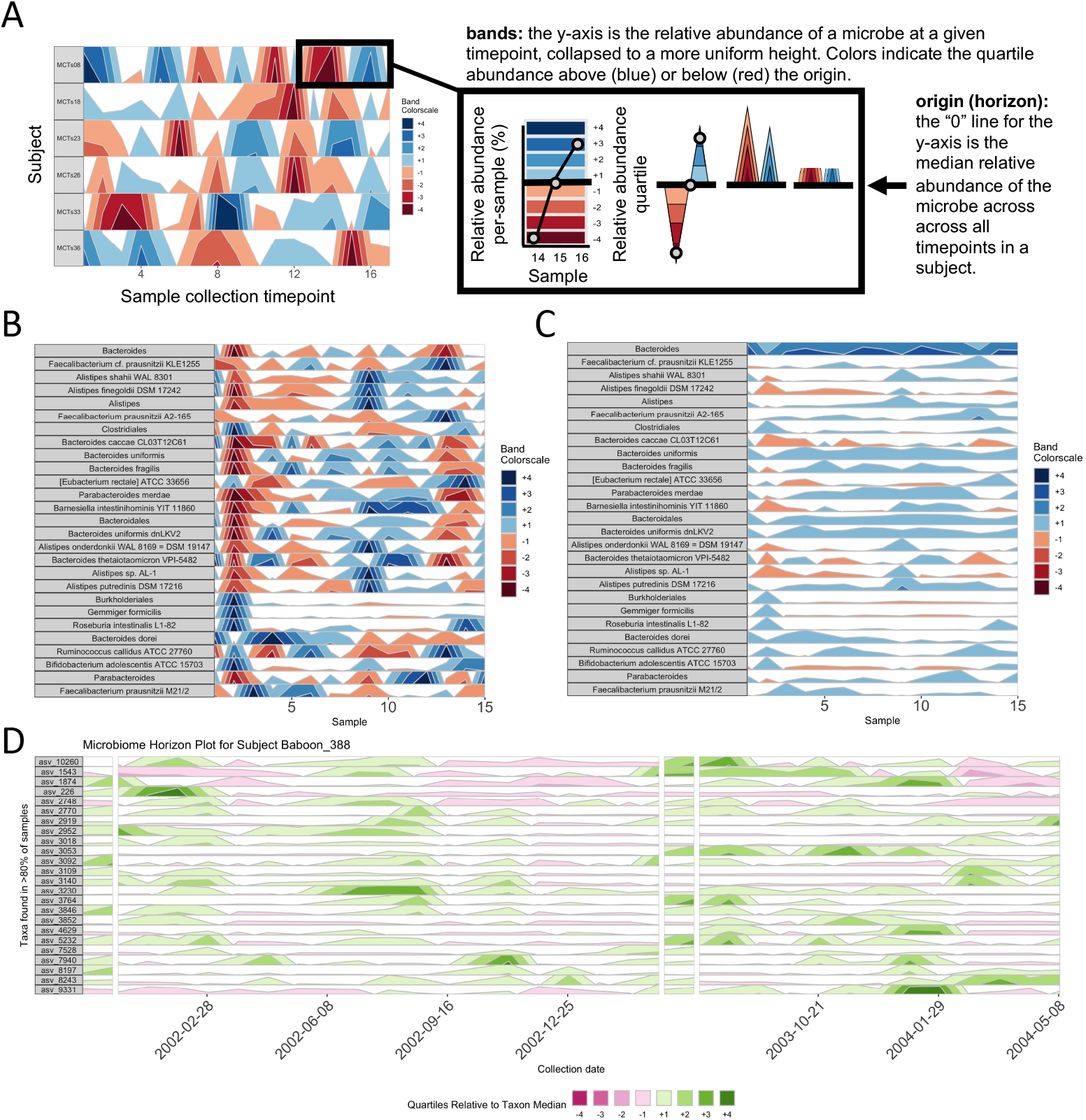
A *BiomeHorizon* horizon plot showing custom configurations. **(A)** Annotated horizon plot for a single microbe (`single_var_otu = ‘Taxon 1’`), for 17 samples across 6 subjects in the diet study example data set. **(B)** Microbes manually chosen as those with a per-sample average relative abundance of at least 0.75% (`thresh_abundance = 0.75`) across 15 samples in one subject (`subj = ‘MCTs01’`) in the diet study example data set. Microbes are labeled by their most fine-grained level of taxonomic identification (`facetLabelsByTaxonomy = TRUE`). **(C)** The same data as (B), but with the origin manually set to 1% relative abundance (origin = 1) and band thickness set so each band represents 10% relative abundance (band.thickness = 10), which serves to visually emphasize changes in highly abundant microbes (e.g. *Bacteroides*) **(D)** For data collected at irregular time intervals or with collection gaps (shown in the wild baboon study example data set), *BiomeHorizon* can interpolate between points to regularize intervals (25 days shown here; `regularInterval = 25`) with breaks when there are gaps greater than a specified interval (75 days shown here; `maxGap = 75`). Custom aesthetics can be used to adjust labels, colors, etc.

*BiomeHorizon* provides three innovations to prior horizon plot applications, which allows substantial room for customization or adaptation to emphasize different aspects of microbiome data (Tan *et al*., 2019; Barandas *et al*., 2020). First, it takes data in a common OTU table format, with functions to filter taxa based on prevalence and abundance. A taxonomy table and metadata table may be supplemented such that microbes can be annotated by taxonomic level and selected from subject(s) of interest, respectively. These datasets are accepted in a wide range of formats, making the package broadly applicable to many experimental and observational conditions. Second, *BiomeHorizon* allows for comparison of dynamics between abundant and rare taxa with scale options, such that *proportional* changes can be seen. Specifically, the tool can switch between a fixed origin or y-scale for each microbe, and variable origin or y-scale, allowing for comparisons between multiple microbes and between multiple hosts. The user can also customize the number of segments data are separated into to differentiate higher values, or to show wider ranges. Third, *BiomeHorizon* can accurately reflect taxa dynamics across irregular time intervals, making it suitable for visualizing data from observational field studies and other datasets with lengthy gaps. These customizations make *BiomeHorizon* versatile in highlighting a wide range of aspects of longitudinal data, and facilitate the user’s ability to pick which microbes may be of interest for additional analyses. We note that *BiomeHorizon* is the first tool for horizon plots specific to microbiome data.

### Usage scenario

To demonstrate the versatility of *BiomeHorizon*, we apply it to two different publicly available microbiome datasets; a 17-day human diet experimental study with metagenomic sequencing of the gut microbiome (Johnson *et al*., 2019), and a multiyear collection of wild baboon 16S rRNA gut microbiome samples (Grieneisen *et al*., 2021). By using `single_var_otu`, *BiomeHorizon* can compare the temporal dynamics of a single microbe across multiple subjects with samples collected on the same days (**Fig. 1A**). Alternatively, by adjusting `thresh_prevalence, thresh_abundance`, or `otulist`, and specifying `subj`, microbes can be filtered by prevalence and abundance, or by name, to compare many microbes in the same subject (**Fig. 1B**). Further, by adjusting `origin, scale`, or `band.thickness`, the dynamics of highly abundant or rare microbes can be emphasized (**Fig. 1C)**. Finally, for data at irregular time intervals, such as those collected in the wild baboon example data set, `regularInterval` specifies the interval at which missing data can be interpolated, while `maxGap` specifies the maximum amount of time between sample collection before a gap in the X axis should be used (**Fig. 1D**).

## Conclusion

*BiomeHorizon* is a powerful tool for visualization of microbiome dynamics over time, as well as a useful initial data exploration tool. It is highly customizable and versatile, as it is designed to accommodate both metagenomics and 16S microbiome data, and can be easily integrated into *ggplot2*, allowing for aesthetic customization. We note it could also be applied to other types of longitudinal data sets (i.e. non-microbiome) that are represented as the relative abundance of many features. *BiomeHorizon* is an open-source project available on GitHub, with a user-guide to supplement the documentation.

## Acknowledgements

We thank members of the Blekhman Lab, especially Sambhawa Priya, for comments on the tutorial. We also thank Abigail Johnson and members of the Amboseli Baboon Research Project, especially Susan Alberts, Beth Archie, and Jenny Tung, for generating the publicly available data sets used in the package. This work was supported by the National Institute of Health [NIH R35-GM12871 to R.B.] and the University of Minnesota Grand Challenges in Biology Postdoctoral Fellowship to L.G.

## Notes

https://github.com/blekhmanlab/biomehorizon/

https://blekhmanlab.github.io/biomehorizon/

### Competing Interest Statement

The authors have declared no competing interest.

## References

Baksi, K.D. et al. (2018) ‘TIME’: A Web Application for Obtaining Insights into Microbial Ecology Using Longitudinal Microbiome Data. Front. Microbiol., 9, 36.

Barandas, M. et al. (2020) TSFEL: Time Series Feature Extraction Library. SoftwareX, 11, 100456.

Costello, E.K. et al. (2012) The application of ecological theory toward an understanding of the human microbiome. Science, 336, 1255–1262.

David, L.A. et al. (2014) Host lifestyle affects human microbiota on daily timescales. Genome Biol., 15, R89.

Flores, G.E. et al. (2014) Temporal variability is a personalized feature of the human microbiome. Genome Biol., 15, 531.

Grieneisen, L. et al. (2021) Gut microbiome heritability is nearly universal but environmentally contingent. Science, 373, 181–186.

Heer, J. et al. (2009) Sizing the horizon: the effects of chart size and layering on the graphical perception of time series visualizations. In, Proceedings of the SIGCHI Conference on Human Factors in Computing Systems. Association for Computing Machinery, New York, NY, USA, pp. 1303–1312.

Human Microbiome Project Consortium (2012) Structure, function and diversity of the healthy human microbiome. Nature, 486, 207–214.

Johnson, A.J. et al. (2019) Daily Sampling Reveals Personalized Diet-Microbiome Associations in Humans. Cell Host Microbe, 25, 789–802.e5.

McKenney, E.A. et al. (2018) The ecosystem services of animal microbiomes. Mol. Ecol., 27, 2164–2172.

Tan, G. et al. (2019) CNEr: A toolkit for exploring extreme noncoding conservation. PLoS Comput. Biol., 15, e1006940.

Turroni, S. et al. (2017) Temporal dynamics of the gut microbiota in people sharing a confined environment, a 520-day ground-based space simulation, MARS500. Microbiome, 5, 39.

Wickham, H. (2016) ggplot2: Elegant Graphics for Data Analysis Springer-Verlag New York.

